# Maturation kinetics of phagosomes depends on their transport switching from actin to microtubule tracks

**DOI:** 10.1101/2022.05.13.491905

**Authors:** Yanqi Yu, Zihan Zhang, Yan Yu

## Abstract

Phagosomes, where internalized pathogens are ingested, mature into a degradative unit through a sequence of dynamical and biochemical changes. The maturation process requires the active transport of phagosomes on actin and microtubules, but how phagosome maturation is kinetically dependent on their actin- and microtubule-mediated transport is poorly characterized. Here, we focused on the early-to-late phagosome maturation and investigated how the kinetic rate of this early maturation process depends on the timing of phagosome transition from actin to microtubules. We performed single-phagosome tracking of their sequential maturation activities, including recruitment of phagosome markers and acidification. Simultaneously, we measured their transport dynamics. We showed that the timing of phagosome transport from actin cortex to microtubules controls the kinetics of early phagosome assembly and transition to late phagosomes. This ultimately determines the fate of phagosome maturation. Our results reveal distinct mechanisms by which the actin- and microtubule-based transport of phagosomes temporally controls the maturation progression of phagosomes during their degradative function.

## Introduction

Phagocytosis is an effective countermeasure exerted by macrophages to internalize and degrade large pathogens. This process begins with the recognition of ligands on the pathogen by receptors on the cell membrane. Immune cells then internalize pathogens into vacuoles called phagosomes and transport them along actin filaments and microtubules from the periphery of the cell to the perinuclear region (1–11). Phagosomes degrade encapsulated pathogens via cascades of biochemical reactions during their intracellular transport. The biochemical reactions include the acidification of the lumen (12), the production of reactive oxygen species (ROS) (13),and the activation of hydrolytic proteases (14). This entire process is known as phagosome maturation. There is a consensus that the intracellular transport of phagosomes along actin filaments and microtubules is indispensable for this maturation process. Importantly, many pathogenic microbes, such as *Salmonella* species, modify the actin and microtubule network of the host cells to subvert endocytic trafficking to keep the integrity of their intracellular vacuole and withstand the degradation of its vacuole by lysosomes (15–18). However, the mechanisms by which the intracellular transport of phagosomes on cytoskeleton track regulates the biochemical activities that occur during their maturation are poorly understood.

So far, studies have been limited to identifying that phagosome transport is involved in the fusion of phagosomes with endosomes and lysosomes. As with endosomes, nascent phagosomes, once formed at the cell periphery, first encounter the actin cortex (1–9, 19). During phagosomal biogenesis, nascent phagosomes continuously fuse with endosomes. This fusion is facilitated by proteins including small GTPase Rab5 and phosphatidylinositol 3-monophosphate (PI3P) (20–24). Phagosome-endosome fusion was found to be attenuated by the disruption of actin or the inhibition of myosin (6, 25, 26). While this indicates that actin-based transport is involved in phagosome-endosome fusion, the exact roles of actin in phagosome maturation are largely unknown. After passing through the actin cortex near the cell periphery, phagosomes are then transported in a bi-directional motion along microtubules, supported by the observation that such transport motion is diminished by microtubule disruption (27). During this microtubule-based transport process, phagosomes fuse with lysosomes to acquire copies of the vacuolar ATPase (V-ATPase), which drives the acidification of phagosomal lumen (28). As early phagosome markers dissociate from the membrane, phagosomes also acquire Rab7, lysosome-associated membrane glycoproteins (LAMPs) and proteolytic enzymes, to transition into late phagosomes and then phagolysosomes for degradation of luminal contents (29). It is believed that microtubule-based transport of phagosomes is required for this process, because microtubule depolymerization leads to a significant decrease in phagosome-lysosome fusion (30), content delivery from late endosomes to phagosomes (31), and acquisition of lysosome markers (32). These conclusions, which were mostly drawn from cytoskeletal disruption experiments, established the important view that the intracellular transport of phagosomes is an integral part of their maturation process. However, findings from those studies do not directly identify the physical mechanisms of how the phagosome transport affects maturation.

Phagosomes must continuously interact with endocytic compartments to complete their maturation biogenesis. The subcellular localization and dynamics of phagosomes and endolysosomes are guided by the polarized tracks of actin filaments and microtubules. Therefore, it is reasonable to hypothesize that the transport of phagosomes on actin filaments and microtubules exerts temporal control over the maturation process by delivering phagosomes and other organelles to the right locations at the right times. However, testing this hypothesis poses unique challenges. Each phagosome functions as an independent signaling unit that evolves over time at its own rate. Additionally, the chemical composition of both its membrane and the contents composition change transiently, as does the dynamics of phagosome transport. Because of such challenges, studies so far have relied heavily on approaches involving pharmacological inhibition to confirm the importance of the phagosome transport without pinpointing the specific mechanisms.

Here, we overcame these challenges by developing particle sensors as phagocytic targets that allowed us to combine dynamic tracking and biochemical imaging of single phagosomes in living cells. With this integrated approach, we identified a distinct mechanism by which the actin- and microtubule-based phagosomes transport regulates the timing of the maturation process. Specifically, we showed that the timing of phagosome transport from the actin cortex to the microtubules controls early phagosome assembly and biogenesis, including the transient recruitment of early phagosome markers (Rab5 and PI3P). This determines the fate and kinetics of the entire phagosome maturation. Taken together, our results elucidate the mechanisms governing the orchestration of the dynamical and biochemical signaling processes that occur during phagosome maturation.

## Result

### Design and characterization of pH-responsive particle sensors

**Figure 1.**
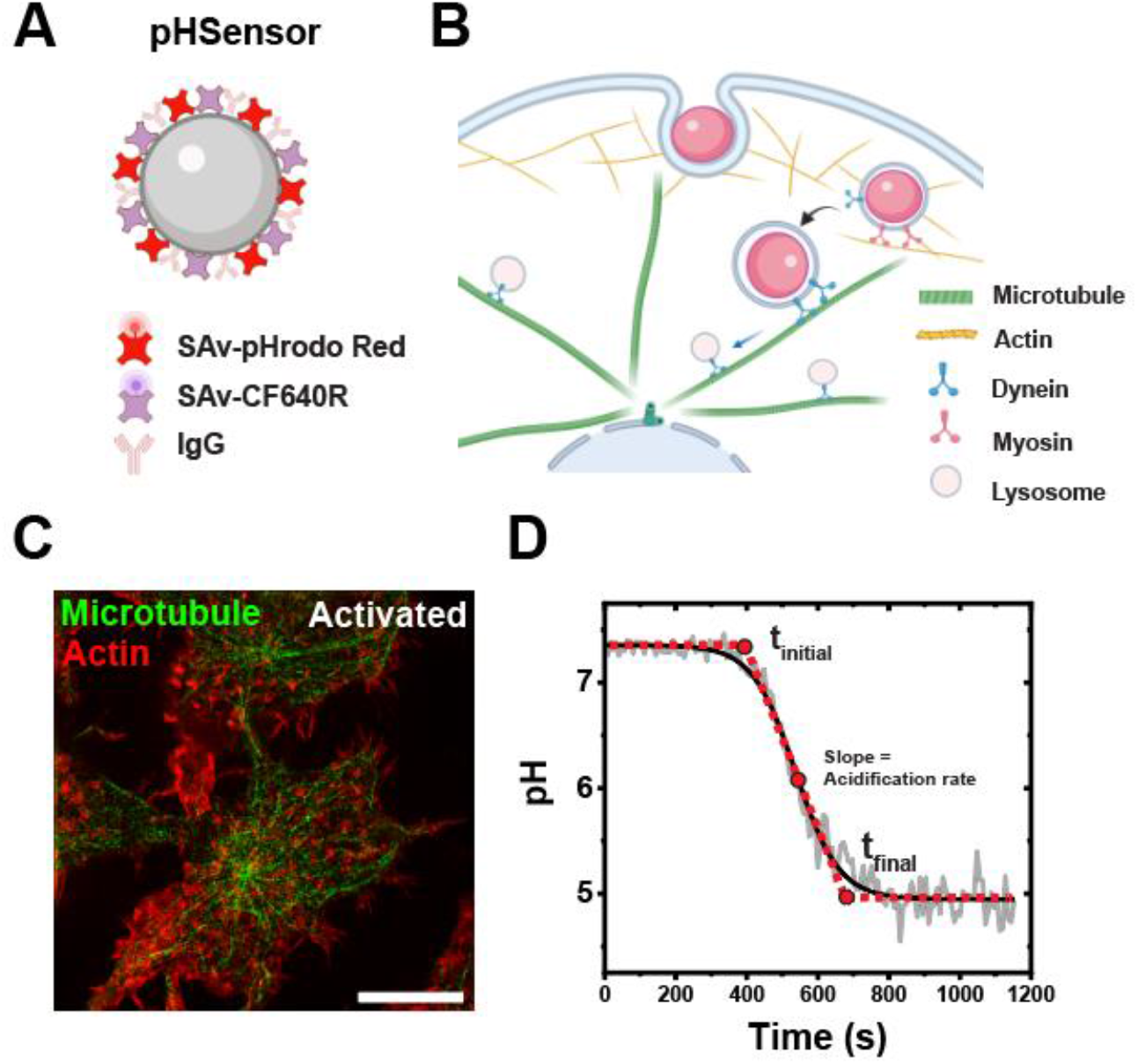
(A) Schematic illustration of the design of pHSensors. Each pHSensors contains a 1-μm silica particles coated with streptavidin (SAv)-pHrodo Red (pH reporter dye) and SAv-CF640R (reference dye). The pHSensors were further opsonized with immunoglobulin G. (B) Schematic illustration of the experimental design. The pH-responsive particle sensors (pHSensors) were internalized into phagosomes in LPS and IFN-γ activated RAW264.7 macrophage cells. (C) Structured illumination microscopy (SIM) images showing immunestained actin (shown in red) and microtubules (green) in LPS and IFN-γ activated RAW264.7 macrophage cells. Scale bars, 10μm. (D) A representative phagosome acidification kinetics reported by pHSensor. The phagosome pH vs. time plot is fitted with sigmoidal-Boltzmann function (shown as the black solid line) to determine the initial pH, final pH, the period of rapid acidification (t*_initial_* to t*_final_*), and acidification rate, as indicated by the red dotted line.

The phagosomal acidification is a key feature of phagosome maturation and a prerequisite for many subsequent maturation events, including the activation of degradative enzymes and optimal processing of antigens for antigen presentation (33, 34). In addition, the intracellular transport of phagosomes appears to be necessary for their maturation, as indicated by arrested acidification and phagosome-lysosome fusion in cells treated with microtubule depolymerization drug nocodazole (30, 35). Therefore, we designed pH-responsive particle sensors (pHSensors) as phagocytic target to measure the both the acidification and intracellular transport of single phagosomes in RAW264.7 macrophages (Figure 1A and B). To fabricate the phagosomal sensors which could faithfully report the pH changes of the phagosome lumen, the 1 μm particle in each pHSensor was biotinylated. Subsequently, pHrodo Red labeled and CF640R labeled streptavidin were allowed to bind on 1 μm particle (Figure 1A). pHrodo Red works as a pH indicator since the fluorescence emission of pHrodo Red increases significantly as the pH of its surroundings decreases from neutral to acidic (36), making it an ideal probe for phagosome acidification. CF640R was chosen as a reference dye because it is insensitive to pH change and photostable.

To quantify the pH sensitivity of the pHSensors, the ratio of fluorescence emission intensities of pHrodo Red and of reference fluorephore CF640R (*I_pHrodc_/I_ref_*,) was measured as we varied pH inside phagosomes in both aqueous buffer and living macrophage cells (Figure S1). For both cases, the ratio *I_pHrodo_/I_ref_* of the RotSensors increased linearly as pH decreases, which is consistent with previous reports (37). When performing the pH calibration, we observed that the pH responses of individual pHSensors, including the initial *I_pHrodc_/I_ref_* ratio and slope, vary slightly from one to another. This was likely due to the different amounts of dyes labeled streptavidin were conjugated on particles and the different lumen environments in phagosomes (38–41). We compensated for the effect of such variation by performing a normalization analysis in all single phagosome pH measurements (details in Materials and Methods section).

We then opsonized pHSensors with immunoglobulin G (IgG) to trigger Fc receptor (FcR)-mediated phagocytosis in classically activated RAW264.7 macrophages (by IFN-γ combined with LPS) (Figure 1C). A representative phagosome acidification profile was shown in Figure 1D. The process started with a standby period where the lumenal pH remained at the extracellular level of ≈ 7.3, and it was followed by a rapid lumenal pH drop to a plateau level at pH 4.5-5.0. Even through individual phagosomes reach slightly different final pH, their pH-time acidification profiles mostly (≈74% of a total of 57 phagosomes) follow a sigmoidal-Boltzmann function:

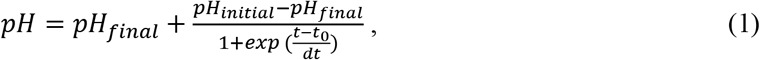

in which t_0_ denotes the half-response point, *t_initial_* and and *tfi_nał_* represent the beginning and the end of the rapid acidification, respectively (Figure 1D). We define the slope at *t_0_* as the acidification rate. The validation of our pHSensors was carried out in cells treated with concanamycin A, a V-ATPase inhibitor. Upon V-ATPase inhibition, the luminal pH of phagosomes stays at neutral value throughout the imaging period (Figure S2).

### Onset time of phagosome acidification depends on actin-to-microtubule based motility transition

**Figure 2.**
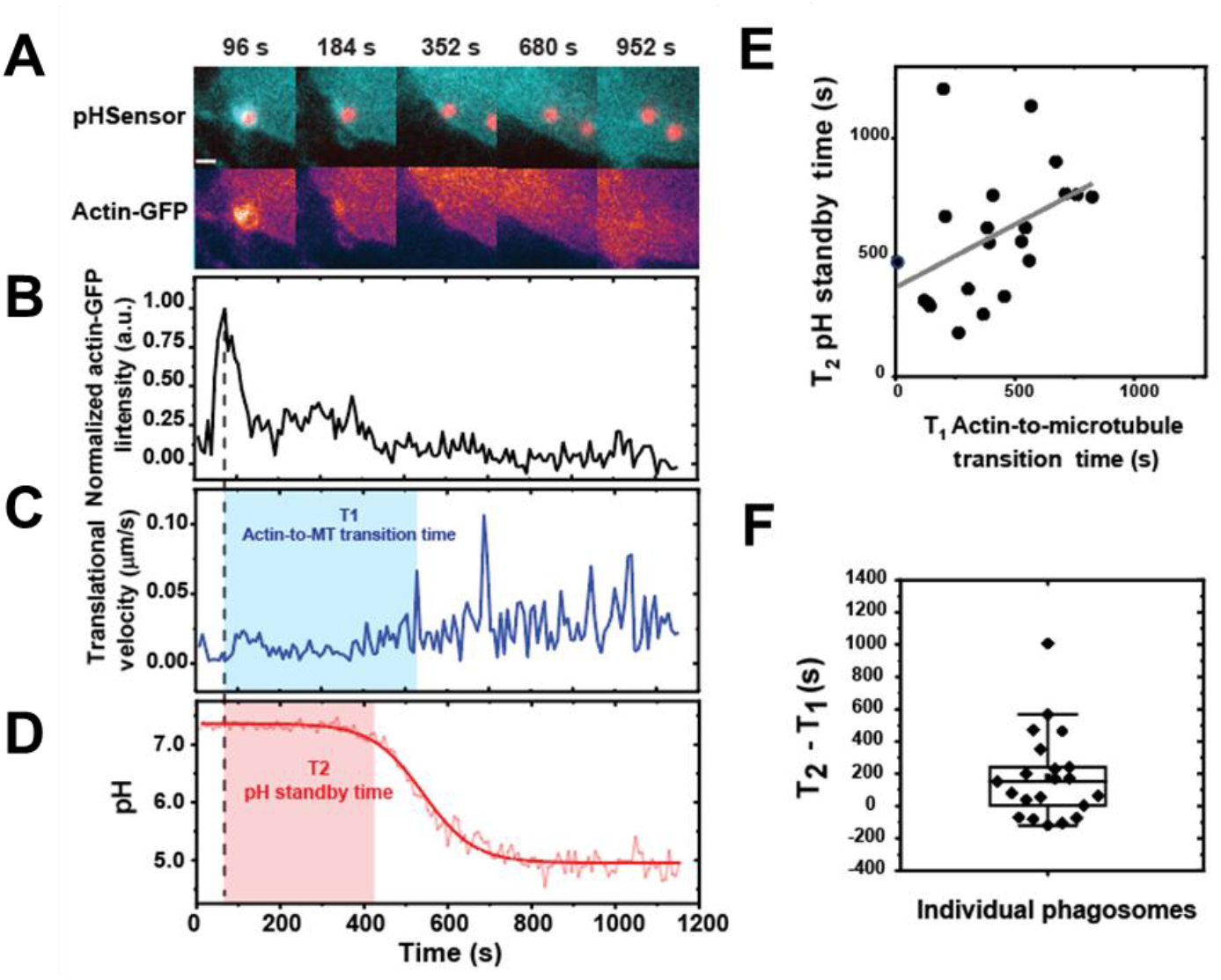
Phagosome dynamics and acidification in LPS and IFN-γ activated actin-GFP expressing RAW264.7 macrophage cells. (A) Fluorescent images showing subcellular locations of pHSensor-containing phagosomes at different time points in actin-GFP expressing cells. Scale bar, 2 μm. (BD) Plots showing the normalized actin-GFP intensity (B), translational velocity (C), and pH (D) of the phagosome shown in (A) as a function of time. In (C), the blue shade indicates the actin-to-microtubule transition period (T_1_). In (D), the thick red line indicates pH after sigmoidal-Boltzmann fitting. The red shade indicates pH standby time (T_2_). (E) Scatter plot showing actin-to-microtubule transition time against pH standby time of single phagosomes in the activated and resting actin-GFP macrophage cells. The thick black line indicates a linear fit to the data points from activated cells, which has a Pearson’s coefficient of 0.44. (F) Box plots showing the time difference between actin-to-microtubule transition time and pH standby time (T_2_-T_1_) for phagosomes in macrophage cells. Each box plot indicates the mean (horizontal line) and the interquartile range from 25% to 75% of the corresponding data set.

Previous studies have shown that intracellular transport of phagosome is composed of distinctive stages with one being actin-dependent transport in cell periphery and the other being microtubule-mediated fast movement towards perinuclear region (2). Here, we aim to study the timing of phagosome acidification and their relationship with distinctive stages of phagosome transport. We imaged phagosome acidification and intracellular transport in classically activated RAW264.7 macrophages expressing actin-GFP, in which intensity changes of actin-GFP can be used to pinpoint the time of particle internalization. As in previous reports (42, 43), we observed that actin rapidly polymerized around nascent phagosomes and then depolymerized at its base, resulting in an actin-GFP intensity peak (Figure 2A and B). This actin intensity peak indicates the completion of engulfment (42). For transport dynamics, we observed that the phagosomes first moved slowly after the internalization, bust later moved bi-directionally and rapidly (Figure 2C). During the slow transport period, their average velocities (0.023 ± 0.015 μm/s) were comparable to that of phagosomes in cells treated with 10 μM of the microtubule inhibitor nocodazole (0.025 ±0.015 μm/s; Fig. S3). This is evidence that the slow movements are independent of microtubules. Using the actin peak as the reference time zero, we found that the rapid microtubule-based transport of phagosomes starts 440 ± 266 sec (mean ± s.d.) after internalization. During this period, phagosomes move from the actin cortex onto microtubules; it is therefore referred to as the “actin-to-microtubule transition time” (Figure 2C). In addition, phagosomes start rapid acidification 620 ± 292 sec (mean ± s.d.) after internalization. This time period is referred to as “pH standby time” (Figure 2D). On average, phagosomes start rapid acidification 186 ± 266 sec (mean ± s.d.) after they move onto microtubules (Figure 2F). When plotting the pH standby time of single phagosomes against their actin-to-microtubule transition time, we observed that phagosomes with longer actin-to-microtubule transition time on average had longer pH standby time, as indicated by the linear fitting with a Pearson’s coefficient of 0.44 (Figure 2E). Our results suggest a temporal correlation between the two distinct stages of phagosome transport with phagosome acidification, where phagosomes largely maintain a neutral pH during their transport within the actin cortex, and rapidly acidify after starting transport on microtubules.

**Figure 3.**
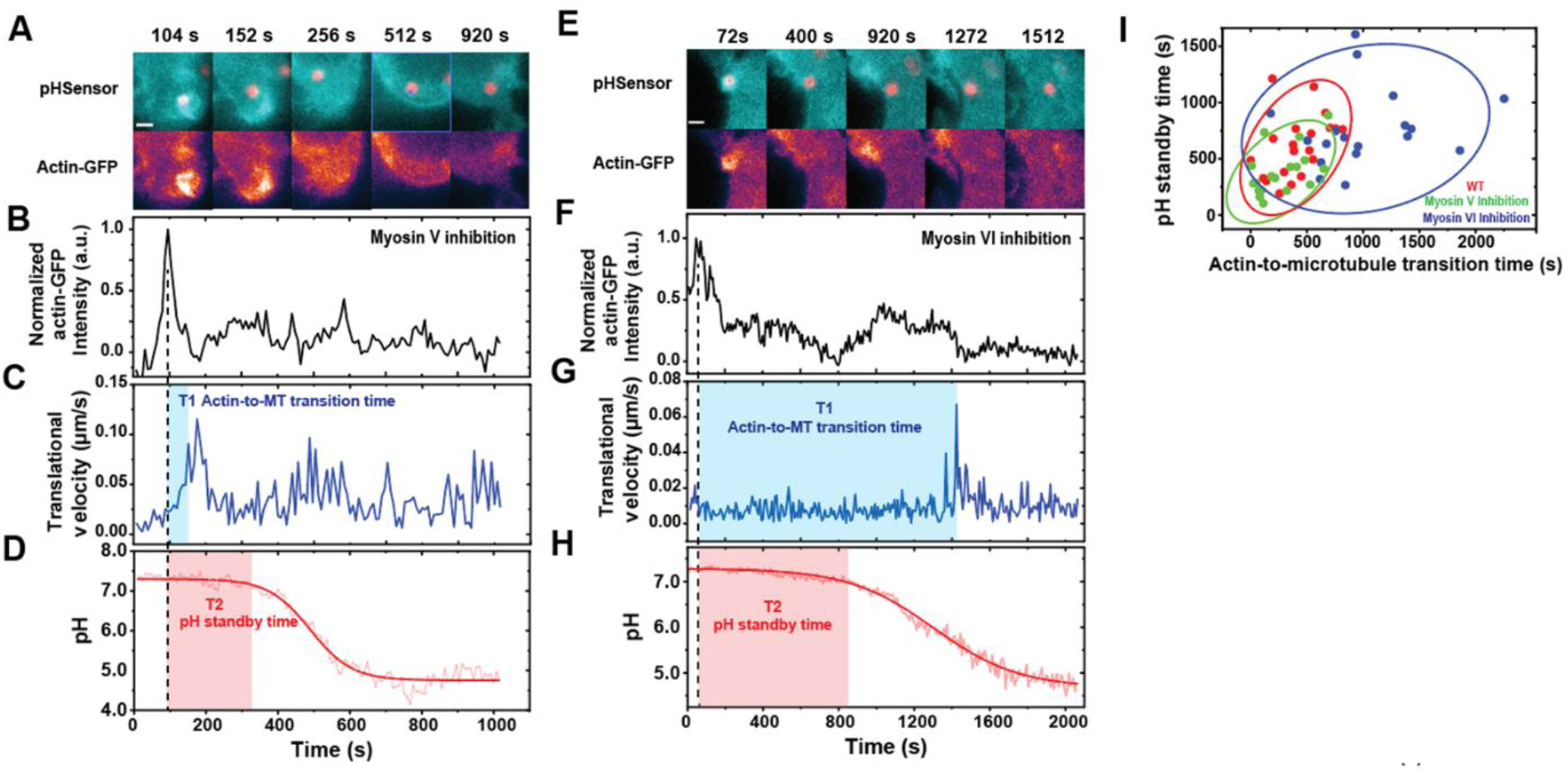
Phagosome dynamics and acidification after Myosin V (A-D) and Myosin VI inhibition (E-H). (A and E) Fluorescence images showing subcellular locations of two single phagosomes at various time points in actin-GFP expressing macrophage cells. Scale bar, 2 μm. (B and F) Plots showing normalized fluorescence intensity of actin-GFP around the phagosome of interest as a function of time. (C and G) Translational velocities of the two phagosomes of interest are plotted as a function of time. The blue shades indicate the actin-to-microtubule transition period (T_1_). (D and H) Plots showing pH vs. time for the two phagosomes of interest. The red shades indicate pH standby time (T_2_). (I) Scatter plot showing pH standby time against actin-to-microtubule transition time of single phagosomes with or without drug treatments as indicated.

To differentiate the roles of the actin- and microtubule-based transport in phagosome acidification, we inhibited Myosin V and VI separately in cells and quantified their effect on the onset time of phagosome acidification. Myosin V moves cargos along actin filaments from the minus end to the plus end, and was shown to tether phagosome and endosomes strongly to the actin cortex and delay their transport onto microtubules (2, 7, 44–46). Contrary to the case of Myosin V, Myosin VI transports cargos from the plus end of actin to the minus end (47). It facilitates cargo transport from actin to microtubules, as its inhibition causes phagosomes and endosomes to be entrapped in actin cortex (8, 48, 49). Indeed, in cells treated with the Myosin V inhibitor MyoVin-1, phagosomes started microtubule-based rapid transport sooner than those in non-treated cells, with a shortened actin-to-microtubule transition time of 276 ±214 sec (Figure 3A-D). On the contrary, phagosomes in cells treated with the Myosin VI inhibitor 2,4,6-triiodophenol (TIP) remained in the actin-based slow-moving stage for a prolonged period of 1026 ± 487 sec (Figure 3E-H). The results confirm that inhibition of Myosin V promotes the handover of phagosomes from actin to microtubules, whereas inhibition of Myosin VI delays this transition. Correspondingly, we found that inhibition of Myosin V led to an earlier onset of rapid acidification (pH-standby time 371 ± 200 sec), and inhibition of Myosin VI led to a delayed onset of rapid acidification (pH-standby time 718 ± 323 sec) compared to non-treated cells (pH-standby time: 620 ± 292 sec). By plotting the pH-standby time of single phagosomes against their actin-to-microtubule transition time (Figure 3I), we confirmed the positive correlation between the two variables. Phagosomes with shorter actin-to-microtubule transition time overall had shorter pH-standby time. It is therefore clear that the timing of phagosome transport from actin to microtubules regulates the onset time of acidification, where phagosomes that moved faster from the actin cortex onto microtubules started acidification sooner.

### Dependence of the rate and final pH of phagosome acidification on actin-to-microtubule based motility transition

**Figure 4.**
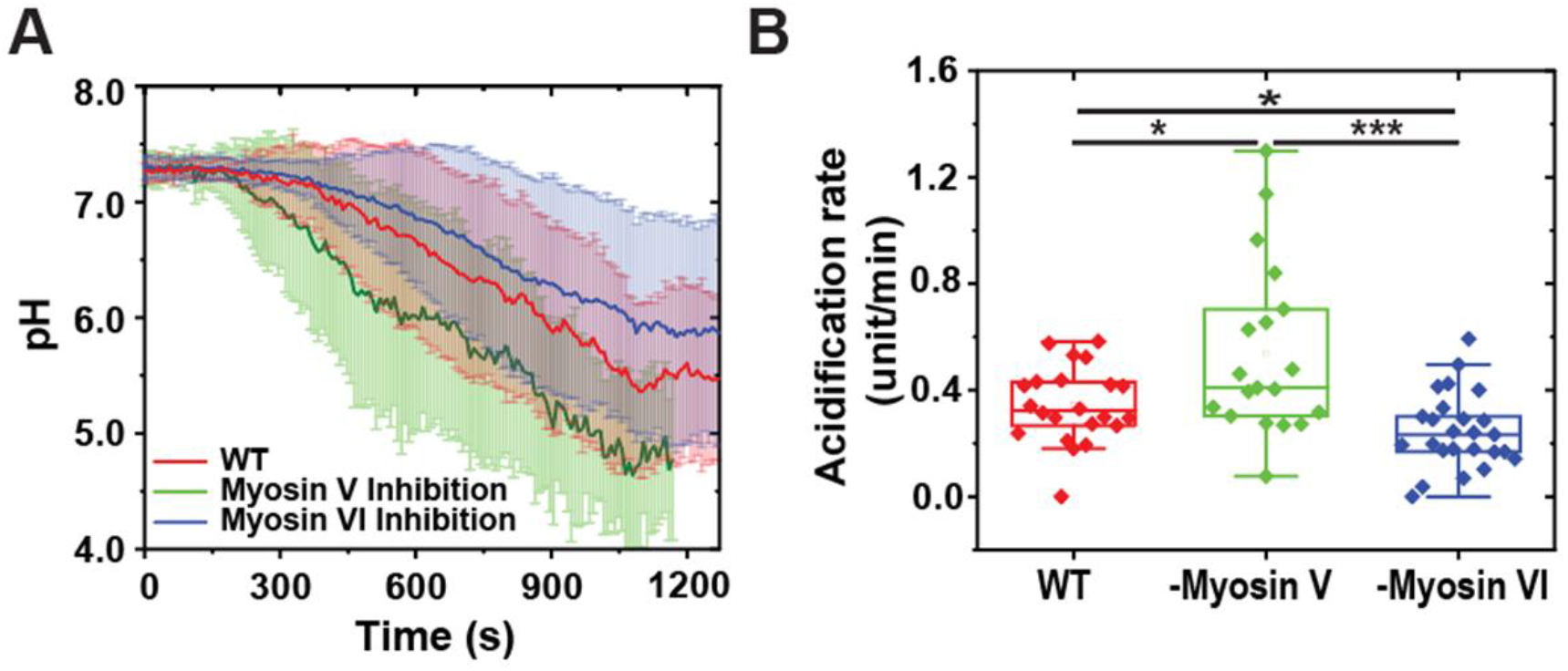
Dependence of phagosome acidification on actin-to-microtubule transport. (A) Line plots showing the average normalized pH as a function of time with or without drug treatments as indicated. The line curves are averaged from 22, 19, and 25 individual phagosomes in cells without drug treatment (wild type), with Myosin V inhibitor, and with Myosin VI inhibitor, respectively. Vertical bars represent standard deviations. (B) Scatter plot showing acidification rate of single phagosomes with or without drug treatments as indicated. The average acidification rate of phagosomes is 0.34 ± 0.14 pH unit/min in cells without drug treatment (WT), 0.54 ± 0.32 pH unit/min with Myosin V inhibition, and 0.25 ± 0.14 pH unit/min with Myosin VI inhibition. In all scatter plots, each box plot indicates the mean (horizontal line) and the interquartile range from 25% to 75% of the corresponding data set. Statistical significance is highlighted by p-values (student’s t-test) as follows: * p < 0.05, ** p < 0.01, *** p < 0.001, NS p > 0.05.

During the analysis of the phagosome acidification upon Myosin V and VI inhibition, we not only observed the dependence of the timing of phagosome acidification on actin-to-microtubule based motility transition, but also the rate of acidification and the final pH of the phagosome lumen that follows (Figure 4A). As shown in Figure 4B, Myosin V inhibition led to a faster acidification (0.54 ± 0.32 unit/min) and a lower final pH (4.5 ± 0.6), compared to non-treated cells (acidification rate: 0.34 ± 0.14 unit/min; final pH: 5.0 ± 0.6). The opposite effect was observed with Myosin VI inhibition: phagosomes exhibited a slower acidification (0.25 ±0.14 unit/min), and a higher final pH (5.5 ±0.8). These observations suggest that actin-based phagosome transport plays have plays an important role in shaping the fate of the maturation.

### Dependence of early-to-late phagosome transition on actin-to-microtubule based motility transition

**Figure 5.**
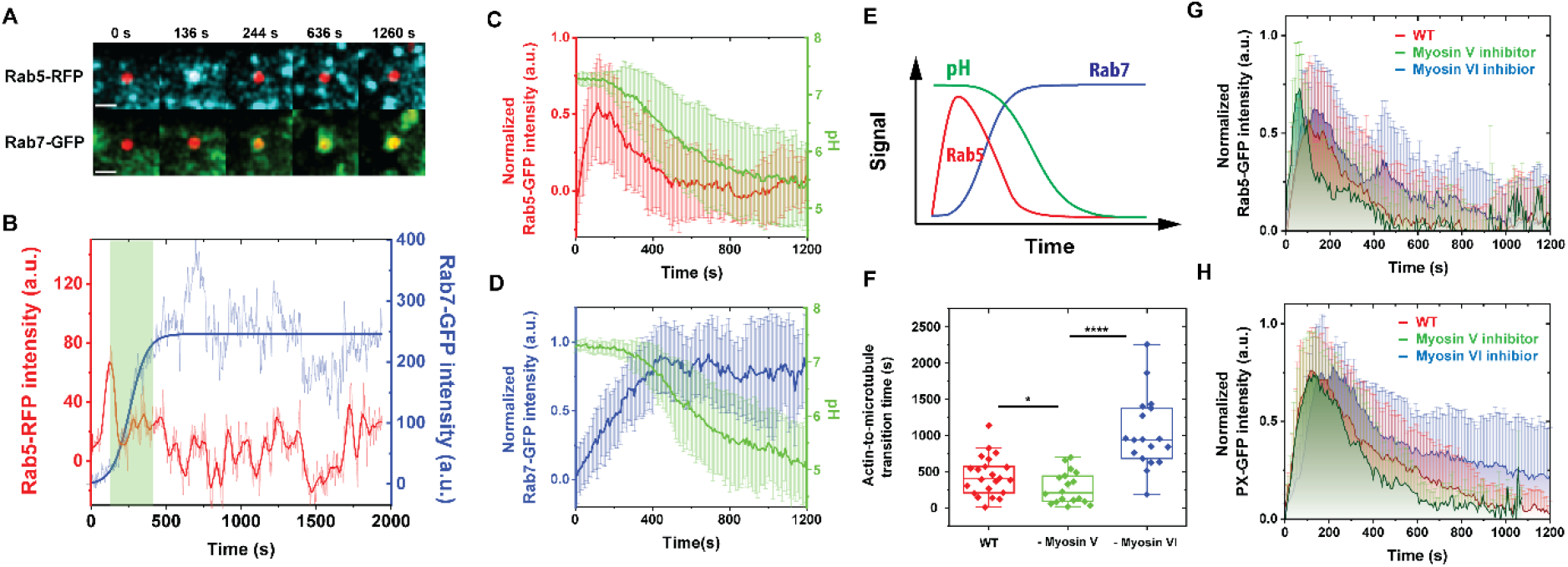
Dependence of early-to-late phagosome transition on actin-to-microtubule transport. (A-B) Fluorescence images and intensity plots showing the dynamic distribution of Rab5-RFP and Rab7-GFP around a representative phagosome during maturation in an activated RAW 264.7 macrophage cell. In the overlaid fluorescence images in (A), the pH-sensor containing phagosome is shown in red, Rab5-RFP in cyan and Rab7-GFP in green. Scale bar, 2 μm. The thick blue line indicates sigmoidal-Boltzmann function fitting to the Rab7-GFP data. The green shaded area indicates the period of Rab5-to-Rab7 transition from the first Rab5 intensity peak to the plateau of Rab7-GFP intensity. (C-D) Line plots showing average normalized intensity of Rab5-GFP (in C), Rab7-GFP (in D), and average normalized phagosome pH (in both plots) as a function of time. The line curves were averaged from 26 and 17 individual phagosomes in Rab5-RFP and Rab7-GFP expressing cells, respectively. Vertical bars indicate standard deviations. (E) A schematic showing the dynamic changes of Rab5, Rab7, and pH during singe phagosome acidification. (F) Scatter plot showing actin-to-microtubule transition time of single phagosomes with or without drug treatments as indicated. The average actin-to-microtubule transition time of phagosomes is 440 ± 266 s in cells without drug treatment (WT), 276 ± 214 s in cells with Myosin V inhibitor, and 1026 ± 487 s in cells with Myosin VI inhibitor. In all scattered plots, each box plot indicates the mean (horizontal line) and the interquartile range from 25% to 75% of the corresponding data set. Statistical significance is highlighted by p-values (student’s t-test) as follows: **** p < 0.0001, * p < 0.05. (G) Plots showing average normalized intensity of Rab5-GFP as a function of time with or without drug treatments as indicated. The line curves were averaged from 26, 10, and 16 individual phagosomes in cells without drug treatment (WT), with Myosin V inhibitor, and with Myosin VI inhibitor, respectively. Vertical bars represent standard deviations. (H) Plots showing the average normalized intensity of p40PX-GFP, a probe for PI3P, in cells under different drug treatment. The line curves are averaged from 12, 14, and 8 individual phagosomes without drug treatment (WT), with Myosin V inhibition, and with Myosin VI inhibition, respectively. Vertical bars represent standard deviations.

We next sought to investigate the role that actin-based phagosome transport plays in the overall process of maturation. Because we observed that phagosomes acidify after transferring onto the microtubules, we expected that the period of phagosome transport in the actin cortex should overlap with early maturation activities prior to acidification. The first maturation activity we measured was the transient recruitment of the small GTPase Rab5 and its subsequent replacement by Rab7. Rab5 and Rab7 are markers for early and late phagosomes, respectively (50, 51). The Rab5-to-Rab7 transition is a key mechanism of the progression of phagosome maturation (29, 50, 52), so we simultaneously monitored both by imaging Rab5-RFP with actin-GFP or Rab7-GFP in cells. We began by confirming that Rab5 was rapidly recruited to phagosomal membrane after particle engulfment, as indicated by the actin-GFP intensity peak (Figure S4). After its transient accumulation, Rab5 intensity gradually decreased over time with large fluctuations. Meanwhile, Rab7 was recruited to the same phagosomes until its intensity reached a plateau (Figure 5A and B). This Rab5-to-Rab7 transition was observed in most phagosomes, but the intensity and duration of the transition varied significantly among different phagosomes (Figure S5A and B). While Rab5 quickly dissociated within a few minutes on some phagosomes, it remained for a prolonged period on others (Figure S6). We then imaged the recruitment of each GTPase in tandem with phagosome acidification (Figure 5C and D). It is clear that the phagosomes remain at a neutral pH when a transiently high level of Rab5 is present on the phagosome membrane. As Rab5 dissociates from phagosomes and Rab7 is recruited, phagosomes start to acidify. The sequence of events is schematically illustrated in Figure 5E. By plotting the recruitment of Rab5 and Rab7 together with the translational velocity of phagosomes as a function of time, we observed that Rab5 recruitment starts during the slow transport of phagosomes within the actin cortex, but its replacement by Rab7 can continue into the period when phagosomes are transported along microtubules (Figure S5 A and B).

To determine the role of actin-based phagosome transport in phagosome maturation, we inhibited Myosin V and VI separately in cells and quantified their effect on the recruitment of Rab5. Correspondingly, we found that the duration of Rab5 association on phagosomes was shortened upon Myosin V inhibition, but prolonged upon Myosin VI inhibition (Figure 5G). We observed the similar effects of myosin inhibition on another early phagosome marker phosphatidylinositol 3-phosphate (PI3P), which is labeled with a genetically encoded biosensor p40PX-GFP. PI3P is a phospholipid generated after Rab5 recruitment by the class III phosphoinositide 3-kinase Vps34 and required for phagosome maturation (53). As shown in Figure 5H, inhibition of Myosin V led to shorter duration of PI3P in the phagosomal membrane, whereas inhibition of Myosin VI caused the opposite effect. The observations from Rab5 and PI3P consistently show that a faster actin-to-microtubule transition with Myosin V inhibition shortens the duration over which Rab5 and PI3P are present on phagosomes. On the other hand, a slower actin-to-microtubule transition due to Myosin VI inhibition prolongs Rab5 and PI3P association with phagosomes. The results evidently show that the timing of transition of phagosomes from actin onto microtubules regulates the turnover of Rab5 and PI3P on phagosomes.

## Discussion

The intracellular trafficking of phagosomes and endolysosomes are guided by the polarized tracks of actin filaments and microtubules. Previous studies have shown that intracellular transport of phagosomes is an integral part of the maturation process (54). However, the physical mechanisms of how the actin-based and microtubule-based transport affects maturation is lacking. In this study, we developed pH-responsive particle sensors as phagocytic targets to simultaneously probe biochemical activity and intracellular transport of single phagosomes in living cells. This approach allowed us to discover a distinct mechanism by which the transport of phagosomes on actin filaments and microtubules exerts temporal control over the maturation process, where phagosome’s transport from actin cortex to microtubules regulates the assembly of proteins to nascent phagosomes and the subsequent maturation progression.

The key finding of our study is that the translocation of the phagosome from cortical actin to microtubule regulates the duration of Rab5 and PI3P recruitment to phagosomes, which subsequently influences the progression to late phagosomes (Rab7 recruitment) and the acidification. A consensus is that nascent phagosomes, once formed, are transported away from the cortical region as they displayed an actively switch from actin-based movement to microtubule-based movement. Importantly, we found that the period of phagosome transport from actin cortex onto microtubules regulates the initiation of phagosome maturation. Previous studies have shown that nascent phagosomes continue to acquire the molecules needed for maturation throughout their transition in the actin cortex (55–57). In particular, proteins such as Rab5 and PI3P, which are required for the fusion of phagosomes with early endosomes (23, 24), are recruited onto phagosome membranes before microtubule-based transport (20–22). However, the role of actin in this process is unclear. Our results here demonstrate that the actin-to-microtubule translocation of phagosomes plays a much more significant role than merely proceeding coordinately with the early-to-late phagosome transition. We showed that prolonging the entrapment of phagosomes in actin cortex through Myosin VI inhibition significantly extended their early phagosome phase. This was indicated by the observation that the duration of Rab5 and PI3P association was prolonged, and the onset of phagosome acidification was delayed. On the other hand, speeding up the transport of a phagosome from actin to microtubule transport through Myosin V inhibition shortened the early phagosome phase and accelerated its transition to a late phagosome. In addition, we found that the final pH of a phagosome is affected by its transport within the actin cortex prior to moving onto microtubules. Our findings reveal that the timing of phagosome transport from actin filaments to microtubules involves a mechanism that directly regulates early phagosome biogenesis, which then, in turn, regulates the subsequent cascades of phagosome maturation activities, including acidification. Several pathways exist by which the duration of phagosome entrapment in the actin cortex might control the assembly of Rab5 and PI3P onto phagosomes. Membrane-bound actin was shown in *in vitro* assays to facilitate the fusion of purified phagosomes with endosomes (58) and between purified yeast vacuoles (59). The disruption of actin was also found to reduce the delivery of early endosomal content to pathogenic mycobacteria containing phagosomes (25). Therefore, the retention of phagosomes in cortical actin might control the timing of their fusion with early endosomes to acquire proteins such as Rab5. Meanwhile, the timing of switching onto microtubules can affect the delivery of Rab5-activating protein 6 (RAP6, also known as Gapex-5) (20), which, in turn, controls the rate of the Rab5-to-Rab7 conversion. Finally, switching onto microtubule promote phagosomes’ fusion with lysosomes (27, 28, 30, 32, 60–63). By fusion with lysosomes, phagosomes acquire proteins essential for maturation, including V-ATPase for acidification (28).

Individual phagosomes, even those from the same cells, exhibit a significant level of variation in maturation kinetics. There is no known mechanism that regulates the individual rate of phagosome maturation. Our results demonstrate that the intracellular transport of phagosomes functions as a maturation clock. Phagosome interaction with the cytoskeleton network can shape their degradative functions. This finding provides new insights into how pathogens might alter the subcellular environment in host cells, including the cytoskeletal structure and activities of molecular motors, to manipulate the degradative capacity of phagosomes as an immune-evasion strategy.

## Materials and Methods

### 1. Materials

Alexa Fluor 488 anti-tubulin-α antibody, Alexa Fluor Plus 647 phalloidin, Pentylamine-biotin, pHrodo iFL Red STP Ester, Pentylamine-biotin, and Streptavidin were purchased from ThermoFisher (Waltham, MA). Immunoglobulin G from rabbit plasma, albumin from bovine serum (BSA), biotin N-hydroxysuccinimide ester (biotin-NHS), nocodazole, MyoVin-1, 2,4,6-triiodophenol (TIP), and lipopolysaccharides were purchased from Sigma-Aldrich (St. Louis, MO). Monodisperse amine-modified silica particles (diameters 1.0 μm) were purchased from Spherotech Inc. (Lake Forest, IL). 1-(3-Dimethylaminopropyl)-3-ethylcarbodiimide hydrochloride (EDC) was purchased from Alfa Aesar (Haverhill, MA). Nigericin sodium salt was purchased from Tocris Bioscience (Minneapolis, MN). Recombinant Murine IFN-γ was purchased from Peprotech (Rocky Hill, NJ). CF640R-amine was purchased from Biotium (Fremont, CA). FuGENE HD transfection reagent was purchased from Promega (Madison, WI). GFP-Rab5, RFP-Rab5, GFP-Rab7, and p40PX-GFP were prepared as previously described (64–66). RAW264.7 macrophage was purchased from ATCC (Manassas, VA). RAW264.7 macrophages stably expressing EGFP-actin have been previously described (42). Ringer’s solution (pH = 7.3, 10 mM HEPES, 10 mM glucose, 155 mM NaCl, 2 mM NaH_2_PO_4_, 5 mM KCl, 2 mM CaCl_2_, 1 mM MgCl_2_) was used for live-cell imaging. Potassium-rich solution (135 mM KCl, 2 mM K2HPO_2_, 1.2 mM CaCl_2_, 0.8 mM MgSO_4_) was used for intracellular pH calibration. Artificial lysosome fluid (55 mM NaCl, 0.5 mM Na_2_HPO_4_, 0.26 mM trisodium citrate dihydrate, 0.79 mM glycine, 150 mM NaOH, 108 mM citric acid, 0.87 mM CaCl_2_·H_2_O, 0.27 mM Na_2_SO_4_, 0.25 mM MgCl2·H_2_O, 0.46 mM disodium tartrate, 1.6 mM sodium pyruvate) was prepared following a previously reported protocol (67) and used for washing particles during the protein conjugation step of the pHSensor fabrication.

### 2. Cell Culture, Pharmacological Treatments and Transfection

RAW264.7 macrophage and the cell lines stably expressing EGFP-actin were cultured in Dulbecco’s Modified Eagle Medium (DMEM) complete medium supplemented with 10% fetal bovine serum (FBS), 100 units/ml penicillin and 100 μg/ml streptomycin at 37°C and 5% CO2. Resting macrophages were activated with a combination of 100 units/ml IFN-γ and 50 ng/ml LPS for 9 hr before using for live cell imaging. For Myosin inhibition, Myosin V inhibitor MyoVin-1 and Myosin 6 inhibitor 2,4,6-triiodophenol (TIP) was separately dissolved in DMSO and added to cells 10 min before live cell imaging. The working concentration was 15 μM for MyoVin-1 and 5 μM for TIP.

For transient transfection, RAW264.7 macrophage cells were seeded on a glass coverslip at 1.0 × 10^5^cells/ml in complete medium for 24 h prior to transfection. Transfection was carried out according to manufacturer’s instructions. In brief, 500 ng of plasmids and 1.5 μl of FuGENE HD transfection reagents were mixed in 100 μl DMEM and kept at room temperature for 15 min. The transfection reagents were then added gently to the cells. Cells were incubated with transfection agents for 24 hr to allow sufficient expression of the encoded protein.

### 3. Fabrication of pH-responsive particle sensors (pHSensors)

#### 3.1 Fluorescence labeling of Streptavidin

Streptavidin-pHrodo Red conjugates (SAv-pHrodo Red) were prepared by mixing 0.7 mg of streptavidin and 60 μg of pHrodo Red STP Ester in 350 μl NaHCO3 solution (100mM, pH 8.2) at room temperature for 3 hr. Unconjugated fluorophores were removed by centrifugal filtration using Amicon Ultra filters (30K). Streptavidin-CF640R conjugates (SAv-CF640) were prepared by mixing 0.7 mg of streptavidin, 80 μg of CF640R-amine, and 3 mg of EDC in 350 μl MES buffer (50mM, pH 4.5) for 3 h at room temperature. Free dyes were removed by centrifugal filtration using Amicon Ultra filters (30K).

#### 3.2 Biotinylation and Streptavidin-biotin Coupling

The first step is the biotinylation of amine-modified non-fluorescent silica particles (1 μm). Amine-modified non-fluorescent silica particles (1 μm) were incubated with 150 μg/ml biotin-NHS in NaHCO3 solution (10 mM, pH 8.25) for 2 hr at room temperature and then with 500 μg/ml of biotin-NHS for 30 min for a second round of biotinylation. After biotinylation, silica particles were washed in methanol and deionized (DI) water, and then incubated in 1×PBS containing 25 μg/ml of SAv-pHrodo Red, 2.5 μg/ml of SAv-CF640R, 5 μg/ml of BSA and 1 μg/ml of IgG for 5 hr at room temperature. Unbound proteins were rinsed off with artificial lysosome fluid and 1×PBS. The resulting pHSensors were further opsonized with 30 μg/ml IgG in 1×PBS for additional 2 hr before adding to cell samples.

### 4. Fluorescence Microscopy

#### 4.1 Single-phagosome pH Assay

RAW264.7 macrophages stably expressing actin-EGFP were seeded on glass coverslips at 2.0 × 10^5^cells/ml in complete medium for 24 h and then activated with a combination of LPS and IFN-γ_following the procedure described above. For all single-phagosome pH studies, the recognition of pHSensors by macrophages was synchronized following a previously reported protocol (68). Briefly, cell samples were placed on ice for 3 min to postpone phagocytosis. pHSensors were then added at a particle-to-cell ratio of ~5:1. Bindings of particles on macrophage were synchronized by centrifuging cell samples at 200×g for 30 s. Live cell imaging was conducted in Ringer’s solution at 37°C on a Nikon Eclipse-Ti inverted microscope equipped with a 1.64 N.A. ×100 TIRF objective (Nikon, Tokyo, Japan) and an Andor iXon3 EMCCD camera (Andor Technology, Belfast, U.K.). Fluorescence emissions at three wavelengths (ex: 488, 561, and 640 nm; em: 515, 586 and 680 nm) were acquired for the time-lapse imaging. The acquisition rate was 8 sec/frame. Single-phagosome pH assays in RAW264.7 macrophages transiently expressing Rab5-GFP, Rab5-RFP, Rab7-GFP and p40PX-GFP were conducted following the same procedure described above.

#### 4.2 Immunofluorescence Staining and Imaging

RAW 264.7 macrophages were first seeded on glass coverslip and then activated as described above. To stain α-tubulin and F-actin, cells were first rinsed with 1× PBS for three times and fixed with 2% PFA at room temperature for 5 min. Subsequently, cell smaples were permeabilized with 0.1% Triton X-100 for 5 min, and blocked with 2% BSA for 1 hr at room temperature. After the blocking, 2 μg/mL Alexa Fluor 488 anti-tubulin-α antibody and 2 μg/mL Alexa Fluor 647 phalloidin were added to cell and allowed to bind for 1 hr at room temperature. Super-resolution structured illumination microscopy (SIM) images of the labeled cells were acquired using the DeltaVision OMX SR imaging system equipped with a Olympus Plan Apo 60×/1.42 PSF objective and a sCMOS camera.

### 5. Images Analysis

#### 5.1 Single-Particle Localization and Intensity Determination

The centroids of single particles in epi-fluorescence images were analyzed using a Gaussian-based localization algorithm in the ImageJ plugin Trackmate. The translational velocity was determined from the centroid location of the 1 μm pHSensor as a function of time. To measure the emission of pHrodo Red and CF640R on pH-Sensors, pixel intensities within a radius of 1 μm from the localized centroid of the particle were integrated and background-corrected using custom MATLAB algorithms. To determine the tracking uncertainties of localization, pHSensors were immobilized on glass coverslip coated with poly-L-lysine and imaged for 200 consecutive frames. Localization uncertainty was defined as the standard deviation of the tracked particle positions in x- and y-coordinates. The localization uncertainty of the 1 μm pHSensor were determined to be 20.6 ± 4.05 nm.

#### 5.2. Intracellular pH Calibration

The intracellular pH calibration of pHSensors was performed following a previously published protocol (38). RAW 264.7 macrophages cultured on glass coverslips were pretreated with 10 μM concanamycin in Ringer’s solution for 10 min. Subsequently, pHSensors were added to and incubated with cells at 37°C for 10 min to promote phagocytosis. After that, DMEM culture medium was replaced with potassium-rich buffers at different pH values. All buffers contain 20 μM of nigericin, but different buffering agent: 5 mM acetic acid for pH 4.5, 5 mM MES for pH 5.5, and 5mM HEPES for pH 6.5 and 7.3. The pH calibration was done from pH 7.3 to 4.5. For each pH, cells were rinsed twice with the correspondingly buffer and allowed to equilibrate for 5 min before image acquisition. Fluorescence emission of the pHrodo Red at 586 nm (*IpHrodoRed*) and reference dye CF640 at 680 nm (I_ref_) was obtained at various pH and background-corrected to obtain ratiometric pH calibration plots (Figure S1). In live cell imaging, pH calibration was performed for individual internalized pHSensors to avoid the effect of particle-to-particle variation in their pH responses. In brief, the ratiometric emission of the pHSensors before acidification was obtained as *I_pHrodo_/I_ref_* at pH 7.3 in each cell sample. After live cell imaging, the same cell sample was incubated in pH 4.5 potassium-rich pH calibration buffer containing 20 μM of nigericin to obtain the pHSensor’s ratiometric emission (*I_pHrodo_/I_ref_*) at pH 4.5. In the final step of the calibration procedure, a linear function was generated based on the known *I_pHrodo_/I_ref_* ratios of the pHSensor at pH 4.5 and 7.3, and the result function was used to derivate luminal pH values from the fluorescence measurements of the pHSensor.

#### 5.3. Image Analysis of the Recruitment of Rab5, Rab7, actin, and PI3P marker

The accumulation of Rab5-GFP around the particle-containing phagosome was analyzed using ImageJ and MATLAB. The following protocol is written using Rab5-GFP as an example, but the same method was applied for analyzing other proteins including Rab5-RFP, actin-GFP, Rab7-GFP and p40PX-GFP. In brief, we first localized the centroid position of the particle in ImageJ. Two circular masks, one with a radius of 1 μm and the other 1.5 μm, were applied and centered at the localized centroid of the particle (Figure S7). The average GFP intensity of all pixels located within the inner circular mask (1 μm radius) was obtained as the signal, and the average fluorescence intensity of all pixels located between the inner and outer circular masks was obtained as the local cytosolic background. The average Rab5-GFP intensity recruited to the phagosome was then obtained by subtracting the cytosolic background from the signal. To compare the duration of Rab5-GFP recruitment between different phagosomes imaged in different cells (as in Figure 5C and Figure S7), we normalized the measured arbitrary intensities of Rab5-GFP to the obtained maximum intensity value in each cell. Same imaging analysis was applied to other proteins including Rab5-RFP, actin-GFP, Rab7-GFP and p40PX-GFP.

## Supplemental information

**Figure S1.**
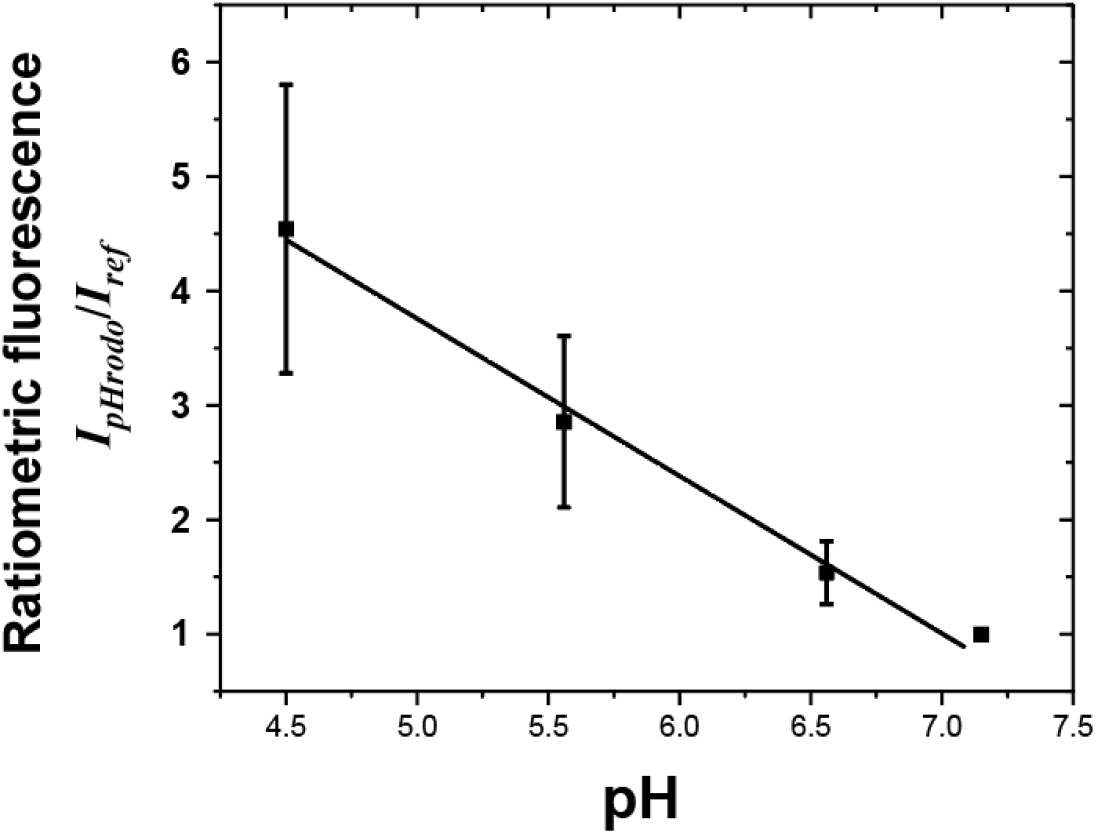
Intracellular calibration of normalized fluorescence emission ratio *I_pHrodo_/I_ref_* of pHSensors (N = 18) as a function of pH. Black line indicates a linear fitting of the data (R2 value of 0.98). The fluorescence emission ratio *I_pHrodo_/I_ref_* at each pH was normalized to that at pH 7.1. Black line indicates a linear fitting of the data (R2 = 0.98).

**Figure S2.**
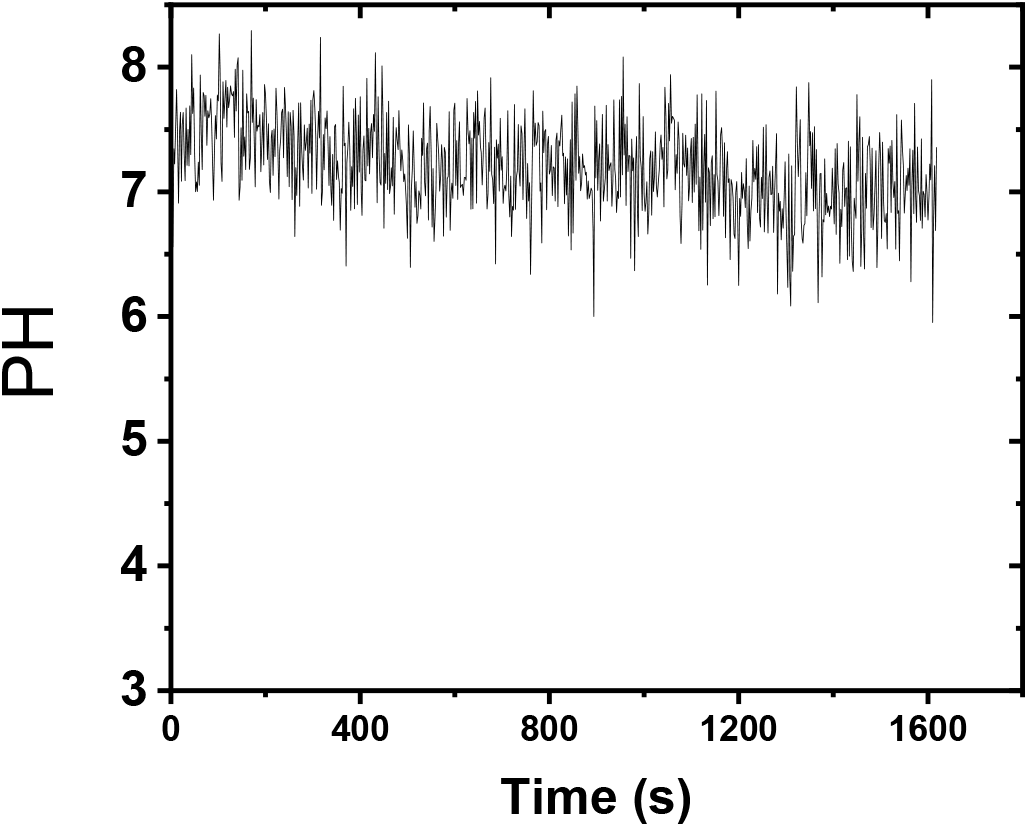
Effect of V-ATPase inhibition on phagosome acidification. Phagosomal pH is reported by pHSensor in macrophages treated with 10 μM concanamycin.

**Figure S3.**
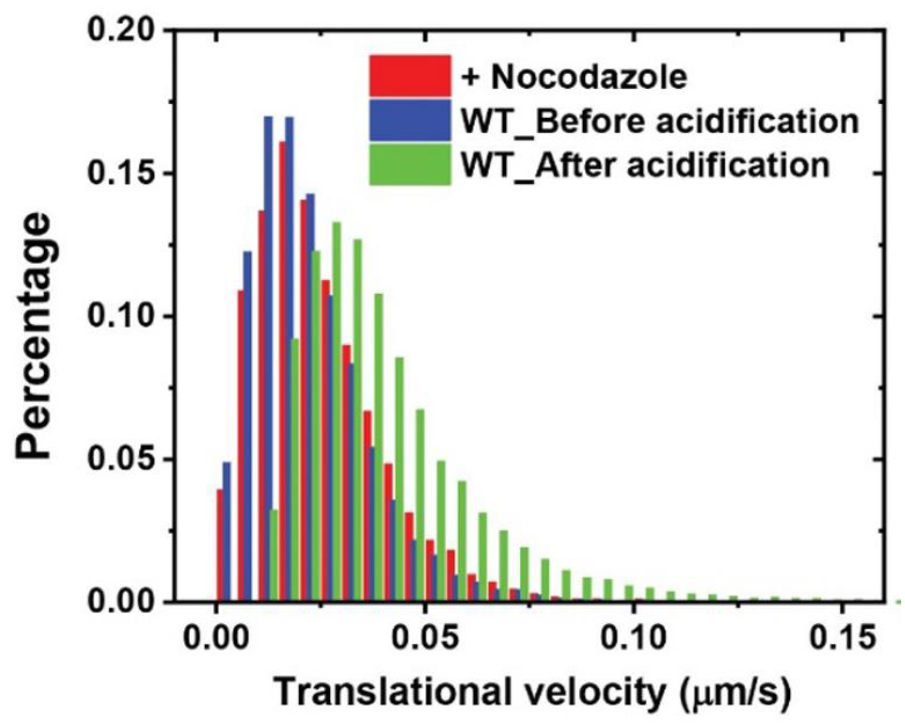
Histograms showing the distribution of (A) translational velocities of phagosomes in control cells (WT) and cells treated with nocodazole, a microtubule-depolymerizing reagent. Data were obtained from 8 individual phagosomes from nocodazole-treated cells and 30 phagosomes from control cells.

**Figure S4.**
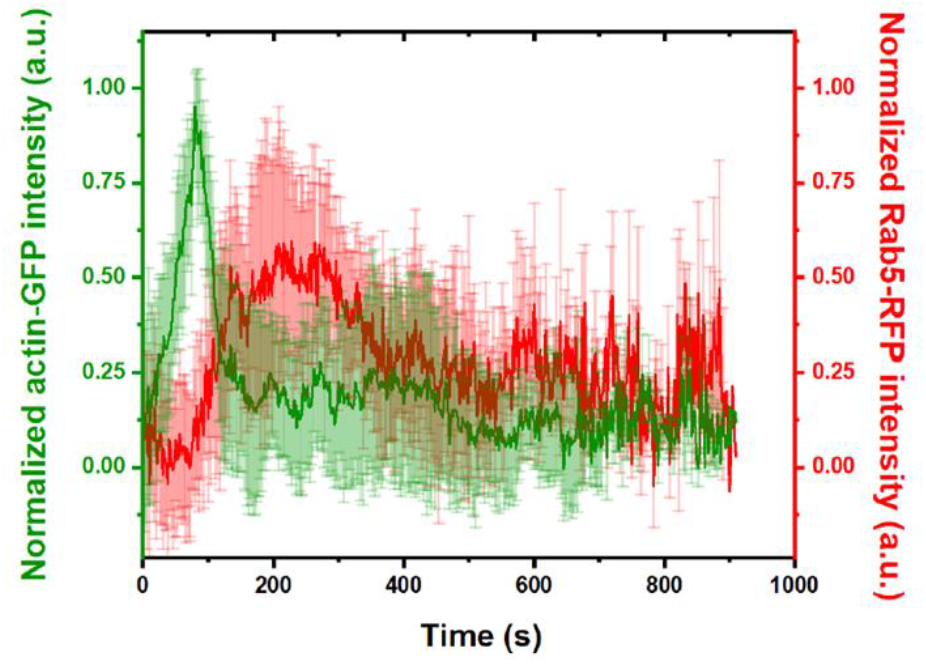
Data showing normalized fluorescence intensity vs. time for actin-GFP and Rab5-RFP on phagosomes obtained from dual-channel simultaneous imaging. Fluorescence intensity of actin-GFP and Rab5-GFP was separately normalized based on its cytosolic background. Each plot is an average of results from 9 independent phagosomes and error bars indicate standard deviations.

**Figure S5.**
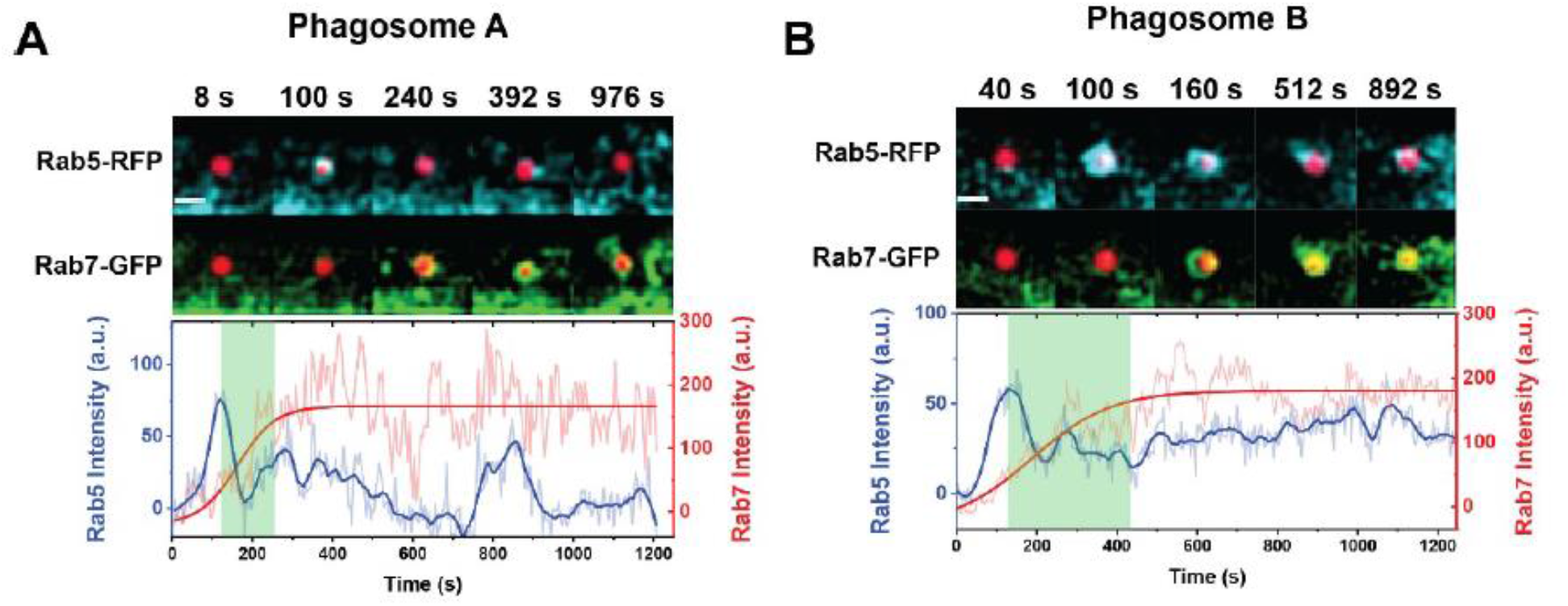
(**A-B**) Fluorescence images and intensity plots showing the dynamic distribution of Rab5-RFP and Rab7-GFP around two phagosomes within the same cell. The fluorescence images show the overlay between pH-sensor (red) with Rab5-GFP (cyan) and Rab7-GFP (green) separately. Scale bars, 2 μm. The green shades in the fluorescence intensity vs. time plots indicate the duration of Rab5-to-Rab7 transition that begins at the Rab5-RFP intensity peak and ends at the onset of Rab7-GFP intensity plateau determined by sigmoidal-Boltzmann function fitting.

**Figure S6.**
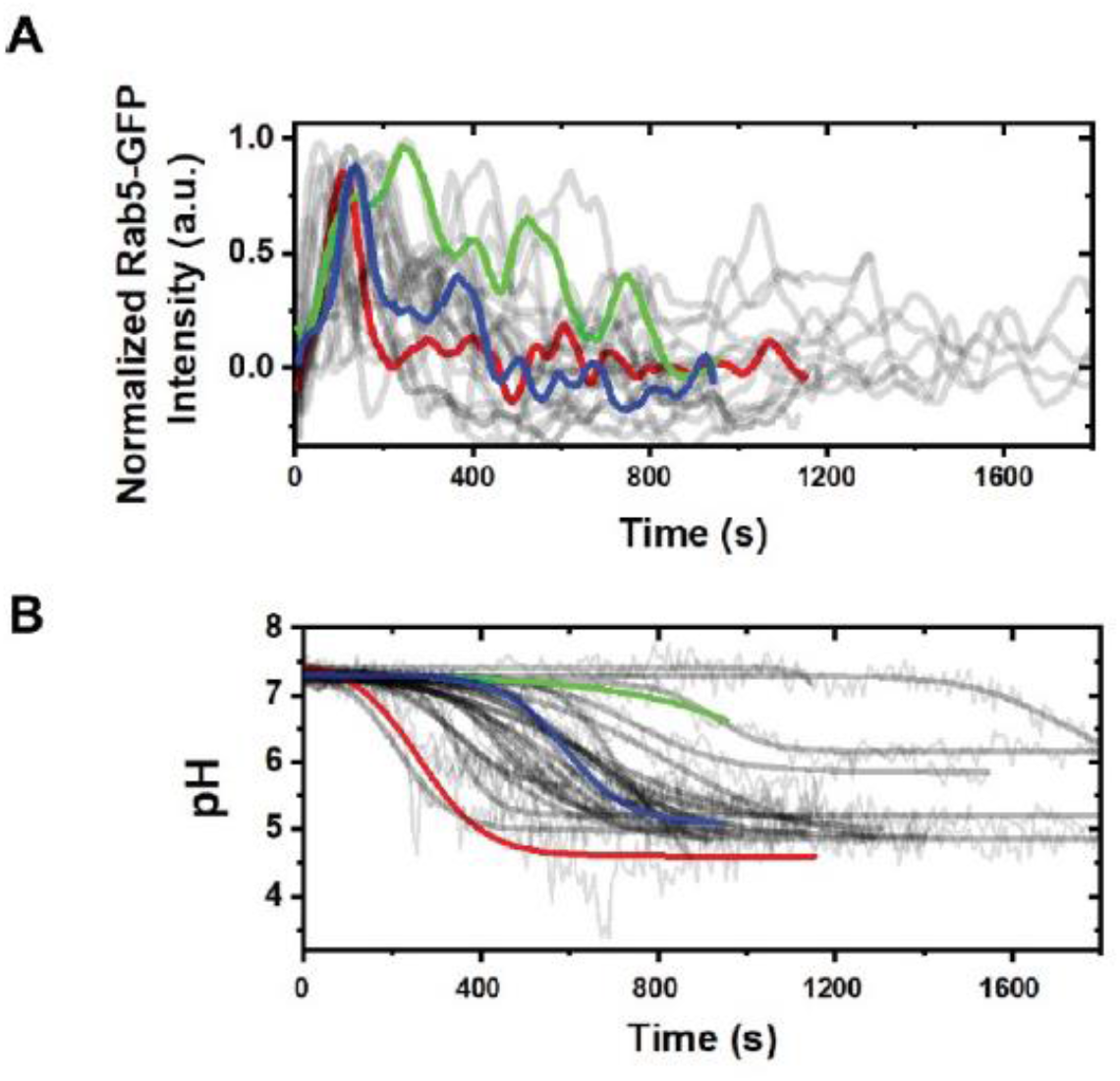
Plots showing normalized fluorescence intensity of Rab5-GFP (**A**) and phagosome pH (**B**) as a function of time. Each curve represents data of a single phagosome. In (**A**) and (**B**), data of three representative phagosomes are highlighted in red, blue, and green. In (**B**), thick lines are sigmoidal Boltzmann function fitting to the individual phagosome acidification profile.

**Figure S7.**
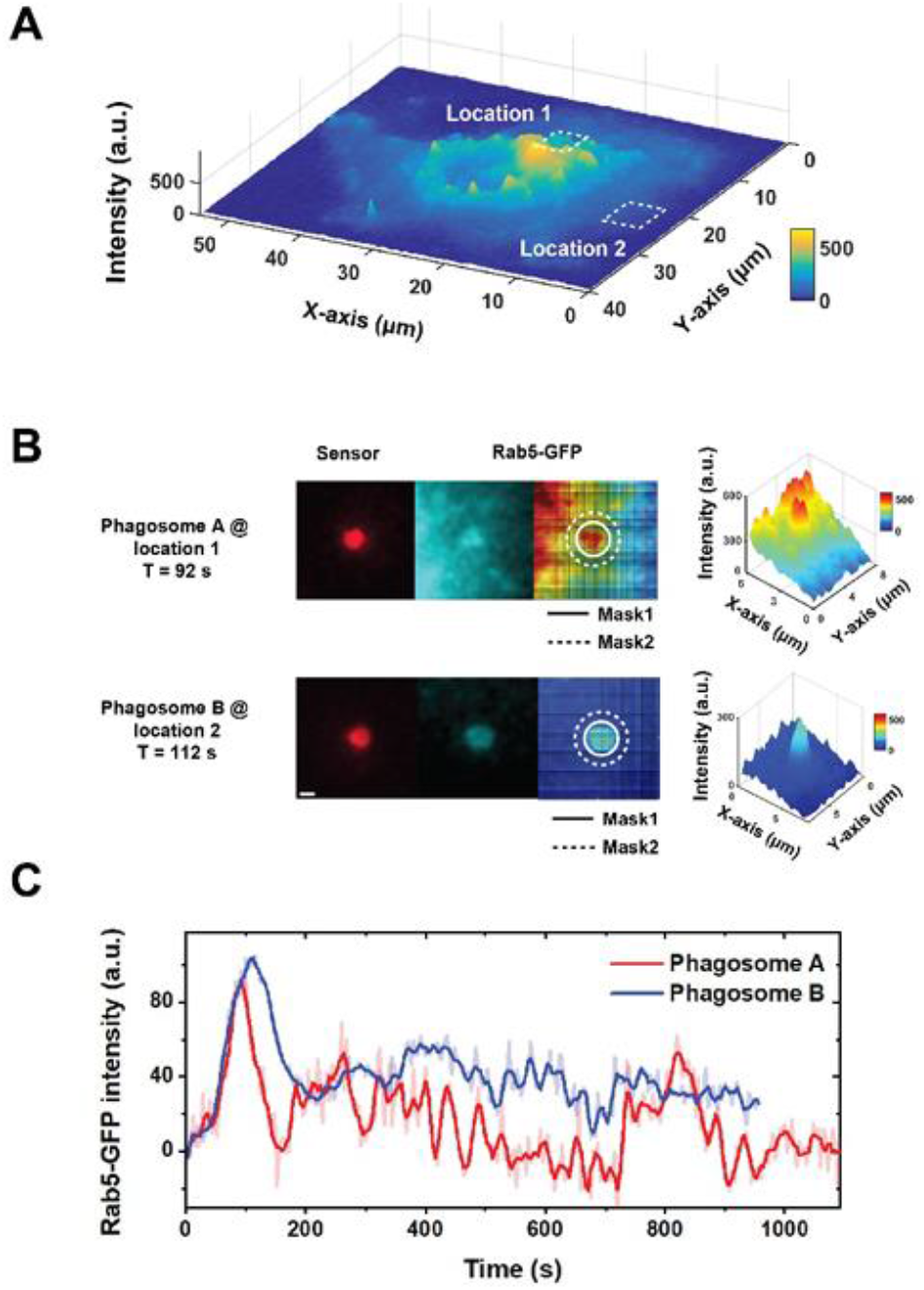
Recruitment of Rab5 on phagosomes differs with subcellular locations. (A) A representative 3D intensity histogram showing inhomogeneous fluorescence distribution of Rab5-GFP in an activated RAW 264.7 macrophage cell (in this case Rab5-GFP). (B) To quantify intensity of Rab5-GFP on single phagosomes and its cytosolic background, two circular masks, indicated as mask 1 and mask 2, were super-imposed on the localized phagosome. The average pixel intensity within circular mask 1 was obtained as signal, and that in between circular mask 1 and 2 was obtained as cytosolic background level. Scale bar, 1 μm. (C) Plots showing the Rab5-GFP intensity after background subtraction for the two phagosomes of interest shown in (A).

